# Understanding the effects of mesenchymal stromal cell therapy for treating osteoarthritis using an *in vitro* co-culture model

**DOI:** 10.1101/2022.08.25.505255

**Authors:** Vivian Shang, Jiarong Li, Christopher B. Little, Jiao Jiao Li

**Affiliations:** Kolling Institute, Faculty of Medicine and Health, University of Sydney, NSW 2065, Australia; School of Biomedical Engineering, Faculty of Engineering and IT, University of Technology Sydney, NSW 2007, Australia

**Keywords:** Mesenchymal stromal cells, Osteoarthritis, Synovial fibroblasts, Co-culture, Inflammation

## Abstract

Osteoarthritis (OA) is a leading cause of chronic pain and disability, for which there is no cure. Mesenchymal stromal cells (MSCs) have been used in clinical trials for treating OA due to their unique functions to send paracrine anti-inflammatory and trophic signals. Interestingly, these studies have shown mainly short-term effects of MSCs in improving pain and joint function, rather than sustained and consistent benefits. This may reflect a change or loss in the therapeutic effects of MSCs after intra-articular injection. This study aimed to unravel the reasons behind the variable efficacy of MSC injections for OA using an *in vitro* co-culture model. Osteoarthritic human synovial fibroblasts (OA-HSFs) exposed to MSCs showed short-term downregulation of pro-inflammatory and pro-catabolic genes, but the MSCs showed upregulation of pro-inflammatory genes and impaired ability to undergo osteogenesis and chondrogenesis in the presence of OA-HSFs. Moreover, short-term exposure of OA-HSFs to MSCs was insufficient for inducing sustained changes to their diseased behaviour. These findings suggest MSCs may not provide long-term effects in correcting the OA joint environment due to adopting the diseased phenotype of the surrounding tissues, which have important implications in the future development of effective stem cell-based OA treatments with long-term therapeutic efficacy.

## INTRODUCTION

Osteoarthritis (OA) is the most prevalent joint disease globally, affecting more than 18% of women and 10% of men over the age of 60 (Glyn-Jones et al., 2015). While OA is generally characterised by the degradation of articular cartilage, disease pathogenesis affects and involves the entire joint including the synovium and subchondral bone (He et al., 2020; Loeser et al., 2012). Due to irreversible damage to joint structures as OA progresses, patients experience joint stiffness, chronic pain, loss of mobility, reduction in quality of life, and an increased risk of mortality from other co-morbidities such as cardiovascular disease (Nüesch et al., 2011). OA occurs in articular joints including the knee, hip, hand, spine, and ankle, with knee OA having the highest prevalence globally (Cross et al., 2014) and estimated to affect more than 650 million individuals aged 40 and over (Cui et al., 2020). The high prevalence of OA and need for chronic disease management due to absence of a cure, places a huge economic burden on healthcare systems worldwide, with direct healthcare costs for OA estimated at 1–2.5% of national gross domestic product in developed countries (Safiri et al., 2020). The global impacts of OA are anticipated to escalate in the coming years due to rising life expectancy, population ageing, and increased obesity prevalence.

Current clinical treatments for OA aim to relieve pain and improve joint functionality in an attempt to delay joint replacement surgery (Cutolo et al., 2015). Non-pharmacologic options focus on patient education, weight loss and exercise programs, but have limited effects on early symptoms and long-term disease modification. Pharmacological treatments generally involve analgesics and non-steroidal anti-inflammatory drugs (NSAIDs), but give rise to inappropriate polypharmacy and increased risk of dangerous side effects due to the high incidence of co-morbidities in OA patients. Intra-articular injections of corticosteroids or visco-supplements such as hyaluronic acid can be indicated for patients whose symptoms cannot be controlled with other treatments. These have yielded variable results, with some evidence supporting short-term effects (less than 6 months) on pain relief and functional improvement (Richards et al., 2016). Surgical intervention is indicated for patients with severe OA who fail to benefit from conservative treatments, where the diseased joint is replaced with an implant to recover joint junction. However, replacement surgery is only available for particular joints (e.g., hip, knee, shoulder) and is associated with increased risks of complications such as infection, and limited implant lifetime of approximately 20 years (Ahmed and Hincke, 2010). The limitations of current OA treatments urge the development of new therapies that may help to stop or delay joint degradation and progression into advanced disease.

Mesenchymal stromal cells (MSCs) have gained attention in OA research due to their potential to provide a disease-modifying and possibly regenerative therapy, owing to their unique immunomodulatory, anti-inflammatory, pro-regenerative, angiogenic, anti-fibrotic, anti-oxidative stress, and multidirectional differentiation functions (Barry, 2019; Harrell et al., 2019). A number of clinical trials have been performed to assess the efficacy of intra-articular MSC injections to treat OA, mostly in the knee (Ha et al., 2019; Jiang et al., 2021). Interestingly, despite most trials reporting short-term symptomatic relief and some regaining of joint function, the therapeutic effects of MSCs diminished within months after administration (Jiang *et al.*, 2021). Some studies have reported that repeated administration of MSCs could help maintain therapeutic efficacy (Matas et al., 2019). It has been proposed that the effectiveness of MSC therapy may be limited by cell survival and engraftment, impeded by the excessive inflammatory immune response, oxidative stress, and hypoxic microenvironments at the injury site (Fernández-Francos et al., 2021). Experiments that traced the engraftment of injected MSCs in OA preclinical models observed that their retention rate was only 3% (Barry, 2019). These findings suggested that the short-term benefits exerted by MSCs in an OA joint were mostly due to their paracrine effects rather than differentiation and replacement of joint cells and tissues, as initially believed.

A wide range of preclinical studies have been performed to elucidate the therapeutic effects and associated mechanisms of MSC activity, in experimental models of OA (Wang et al., 2022; Xing et al., 2018). While the field has built up ample evidence on the benefits of MSC injections in various OA animal models, only one-way analyses were performed in these studies. Since injected MSCs cannot be retrieved for analysis, all *in vivo* findings focus on the effects of MSCs on OA joint tissues, and not the other way around. Furthermore, the diversity of OA disease phenotypes and associated pathophysiology (endotype) in patients cannot be replicated by single animal models (Hunter and Little, 2016; Zaki et al., 2022), which likely contributes to the poor translation of long-term disease modifying therapeutic benefits of MSC injections observed in pre-clinical research to clinical studies, that mostly show only short-term effects.

Very limited evidence exists for exploring the effects of an OA joint environment on MSCs. In one of the only studies that established an “*in vitro* OA cell model”, MSCs were co-cultured with macrophages and OA chondrocytes in “high” and “low” inflammatory environments to simulate different OA joint conditions (Diaz-Rodriguez et al., 2019). Variable anti-inflammatory and immunomodulatory effects of MSCs were observed on the chondrocytes and macrophages depending on the inflammatory state of the culture environment. Coupled with other studies demonstrating that the exposure of MSCs to inflammatory cytokines such as tumour necrosis factor (TNF)-α can lead to increased expression of inflammatory markers in the MSC secretome (Lee et al., 2010), we hypothesise that the inhibitory environment created by diseased cells in an OA joint environment may be at least in part responsible for the reduced therapeutic benefits of MSCs seen in OA clinical studies.

While a range of studies have investigated the effects of MSCs on *in vitro* and *in vivo* OA models, there is an important knowledge gap with regard to: (1) the effects of OA joint-tissue cells on MSCs, and (2) whether a short-term exposure of OA cells to MSCs (similar to the time period of MSC retention after injection into an OA joint) is sufficient for reducing their diseased characteristics in a sustained manner, and thus might improve their response to repeated MSC injections. To answer these questions, we established an *in vitro* co-culture model using human MSCs and primary osteoarthritic human synovial fibroblasts (OA-HSFs). We chose OA-HSFs as the cell model due to their known involvement and contribution to OA pathogenesis (Li et al., 2021), displaying the hallmarks of OA-related inflammation and synovial fibrosis (Maglaviceanu et al., 2021), as well as relative ease of extraction and expansion while retaining diseased characteristics compared to other relevant cell types such as OA chondrocytes (Jackson et al., 2014; Smith et al., 2013). Our findings suggested that the behaviour and therapeutic benefits of MSCs are significantly affected by the OA joint environment, which have important implications for developing effective stem cell-based OA treatments.

## RESULTS

The reciprocal effects of human osteoarthritic synovial cells and MSCs on each other were investigated through short- and long-term co-culture to simulate the types of interactions present in an osteoarthritic joint following stem cell injection. Three co-culture studies were performed to investigate: (1) The effects of MSCs on OA-HSFs in growth medium; (2) The effects of OA-HSFs on MSCs in growth, osteogenic and chondrogenic media; (3) The effects of pre-conditioning OA-HSFs with MSCs prior to co-culturing to mimic repeated injections of MSCs, in growth, osteogenic and chondrogenic media.

### OA-HSFs co-cultured with MSCs rapidly reduced pro-inflammatory and pro-catabolic molecule expression in the short-term

Expression of pro-inflammatory and pro-catabolic molecules implicated in OA pathogenesis was analysed for OA-HSFs cultured in growth medium, with or without MSCs in co-culture (Figure 1). At 3 days, the OA-HSFs co-cultured with MSCs rapidly reduced the expression of several molecules, particularly ADAMTS-5, MMP-13, and TLR4, compared to the control group (Figure 1A). Interestingly at 7 days, there were no differences in expression levels between OA-HSFs which were or were not co-cultured with MSCs for all inflammatory markers except TLR4 and COX2 (Figure 1B). The level of differences between groups also decreased at 7 days compared to at 3 days for all genes. These findings suggest that the MSCs have paracrine anti-inflammatory and anti-catabolic effects on the OA-HSFs, but these effects appear to only persist in the short-term and are not retained in the OA-HSFs.

**Figure 1.**
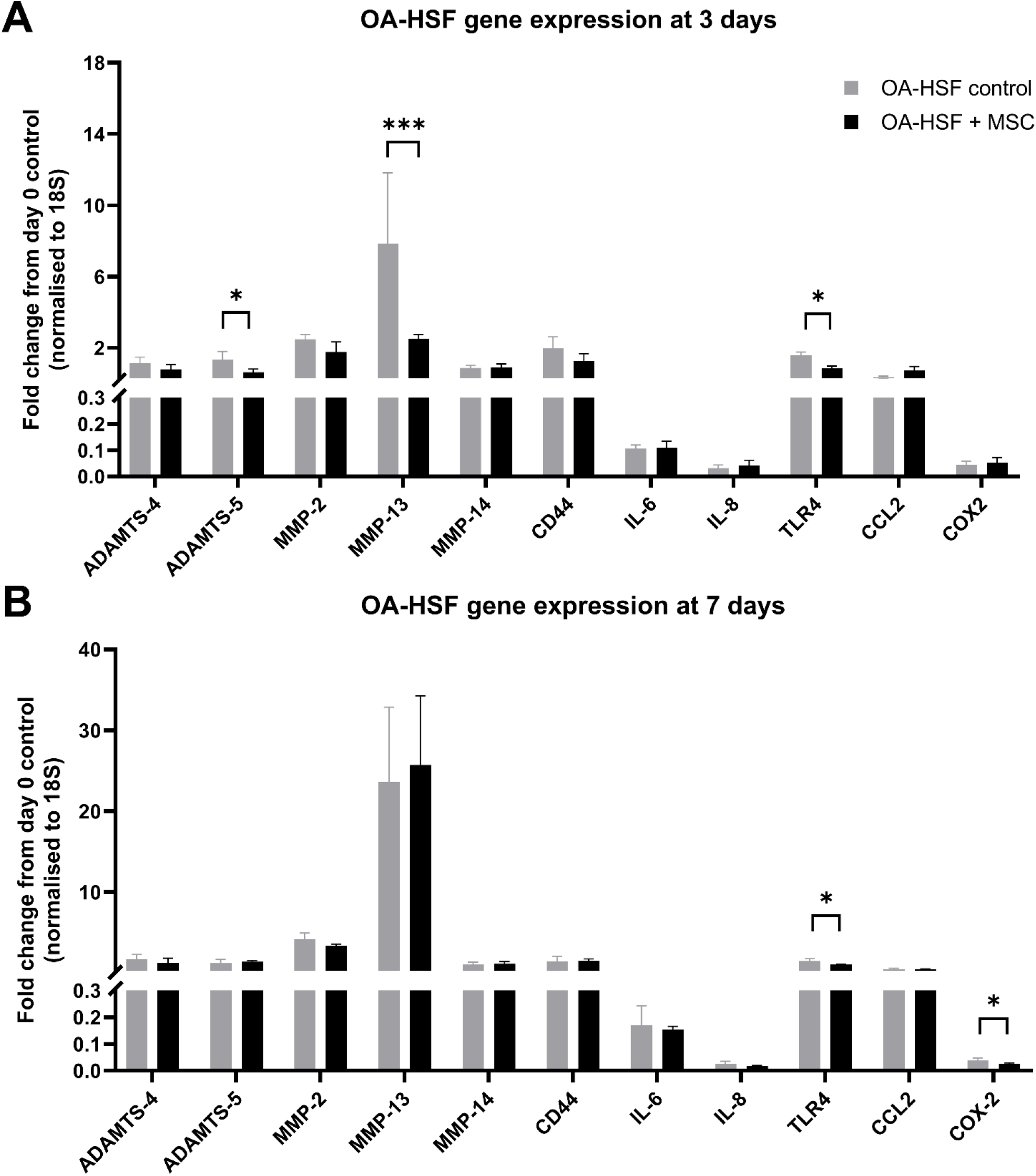
Gene expression of OA pro-inflammatory and pro-catabolic molecules in OA-HSFs. OA-HSFs were cultured in growth medium with or without MSCs in co-culture at **(A)** 3 days, and **(B)** 7 days. Expression levels were presented as fold change from OA-HSFs at day 0, normalised to the housekeeping gene 18S. Data are represented as mean ± SD; N=4; *p < 0.05, **p < 0.01, ***p < 0.001.

### MSCs co-cultured with OA-HSFs showed increased expression of pro-inflammatory and pro-catabolic molecules and impaired differentiation over the long-term

MSCs were cultured in growth, osteogenic, and chondrogenic media to mimic the types of conditions relevant for joint repair. Gene expression of pro-inflammatory and pro-catabolic molecules implicated in OA pathogenesis was analysed for MSCs grown with or without OA-HSFs in co-culture over 21 days (Figure 2). Gene expression and histological staining relevant to osteogenesis (in osteogenic medium) and chondrogenesis (in chondrogenic medium) were also analysed at 21 days for MSCs grown with or without OA-HSFs in co-culture (Figure 3).

**Figure 2.**
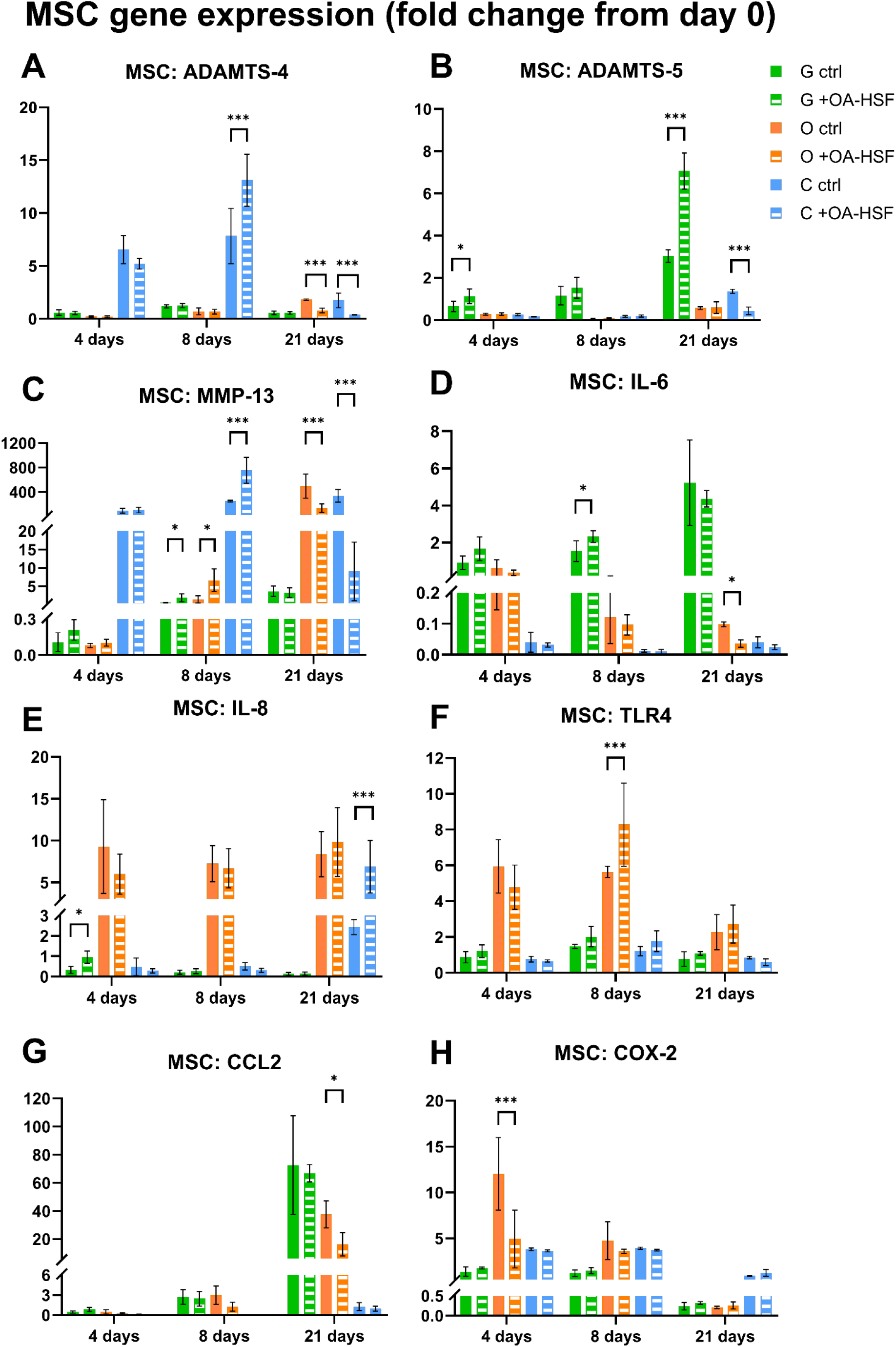
Gene expression of OA pro-inflammatory and pro-catabolic molecules in MSCs. MSCs were cultured in growth (G), osteogenic (O), and chondrogenic (C) media, with or without OA-HSFs in co-culture for up to 21 days. Expression levels were presented as fold change from MSCs at day 0, normalised to the housekeeping gene 18S. Data are represented as mean ± SD; N=4; *p < 0.05, **p < 0.01, ***p < 0.001.

**Figure 3.**
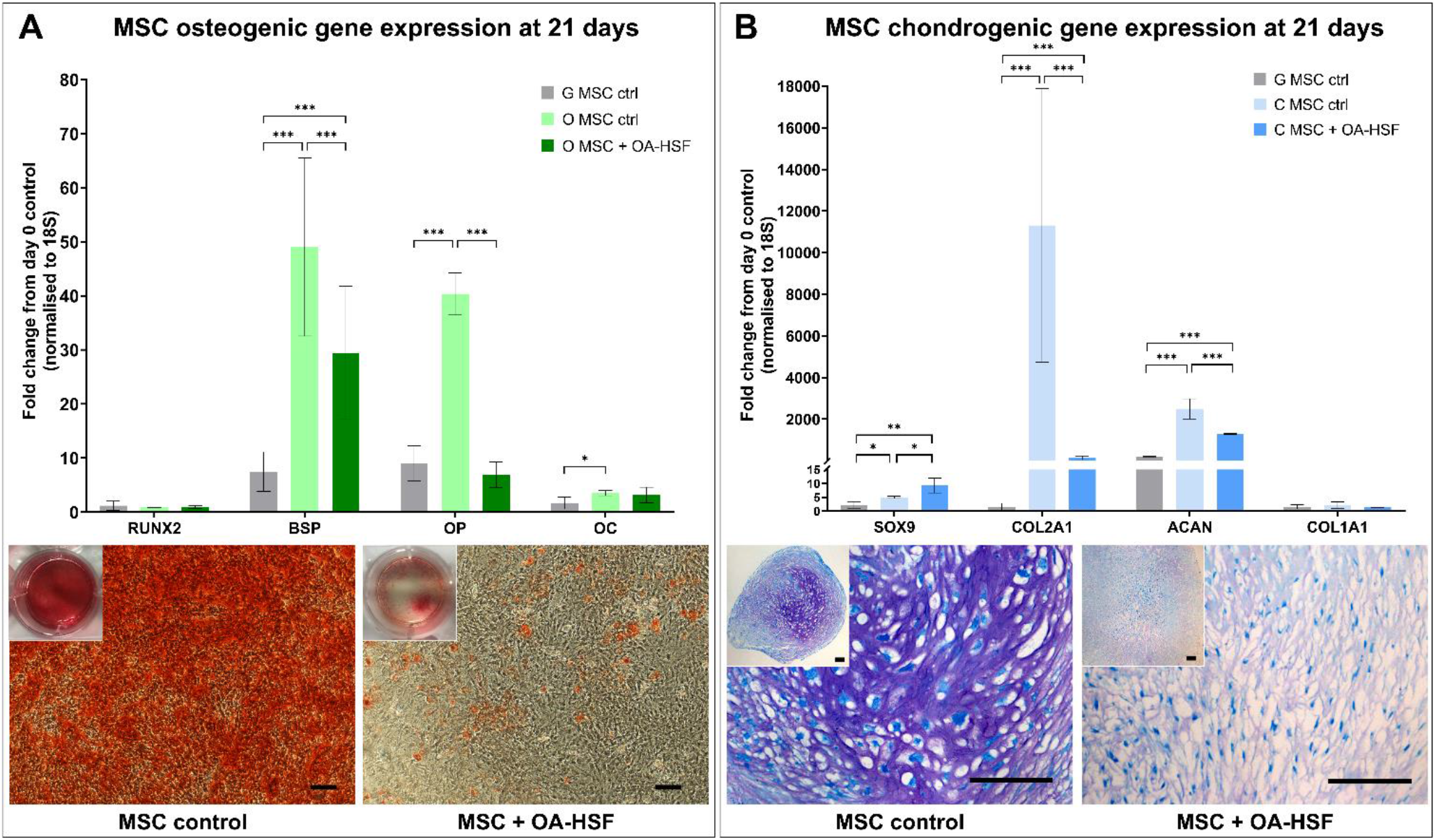
Gene expression and histology of MSCs undergoing differentiation. **(A)** Osteogenic and **(B)** chondrogenic differentiation of MSCs when cultured in osteogenic (O) or chondrogenic (C) differentiation medium, compared to control growth (G) medium, with or without OA-HSFs in co-culture at 21 days. Gene expression levels were presented as fold change from MSCs at day 0, normalised to the housekeeping gene 18S. Data are represented as mean ± SD; N=4; *p < 0.05, **p < 0.01, ***p < 0.001. Calcium deposition in osteogenic cultures is indicated by Alizarin red S staining (3A; scale bar = 500 μm). Proteoglycan deposition in chondrogenic cultures is indicated by toluidine blue staining (3B; scale bar = 200 μm).

A number of genes involved in inflammation and matrix/tissue degradation were found to be elevated in MSCs co-cultured with OA-HSFs compared to MSC controls, in different media types and at both short- and long-term time points. Notably at 8 days, MSCs co-cultured with OA-HSFs showed significantly higher expression levels of MMP-13 in all three media types. Other genes such as ADAMTS-4, ADAMTS-5, IL-6, IL-8 and TLR4 were significantly more highly expressed in MSCs co-cultured with OA-HSFs at 4 or 8 days in at least one media type. For some genes, their upregulation in MSCs co-cultured with OA-HSFs persisted until 21 days, notably for ADAMTS-5 in growth medium and IL-8 in chondrogenic medium. These findings suggest that osteoarthritic cells may create an environment that acts on MSCs and increases their expression of pro-inflammatory and pro-catabolic factors, which may dampen the therapeutic benefits of MSCs. Interestingly, there were a number of inflammatory proteins with reduced expression at 21 days in MSCs co-cultured with OA-HSFs compared to controls, including ADAMTS-4, ADAMTS-5, MMP-13, IL-6, and CCL-2, but exclusively in osteogenic or chondrogenic media, which may be associated with global changes in MSC gene expression at the later stages of differentiation.

MSCs co-cultured with OA-HSFs showed impaired ability to undergo osteogenic and chondrogenic differentiation at 21 days (Figure 3). This was evidenced by markedly reduced gene expression of middle-late stage markers for osteogenesis (BSP, SPP1) and chondrogenesis (COL2A1, ACAN) compared to control MSCs in differentiation medium. MSCs co-cultured with OA-HSFs also showed greatly attenuated histological features of differentiated bone (calcium deposition; Figure 3A) and cartilage (proteoglycan deposition; Figure 3B). These findings suggest that long-term exposure of MSCs to a diseased osteoarthritic environment may cause them to lose their defining characteristics, such as differentiation potential, and result in impaired regenerative ability.

### Repeated exposure of OA-HSFs to MSCs does not change their inflammatory gene expression or influence on MSCs

MSCs were co-cultured with OA-HSFs in growth, osteogenic, and chondrogenic media. The OA-HSFs were either ‘pre-conditioned’ (cHSF groups) or not pre-conditioned (OA-HSF groups). The pre-conditioned OA-HSFs were pre-exposed to MSCs in co-culture for 3 days in growth medium, and subsequently co-cultured with fresh MSCs, simulating repeated injections of MSCs. The OA-HSFs not pre-conditioned were simply grown in growth medium for 3 days before being co-cultured with fresh MSCs. Gene expression of proteins implicated in OA pathogenesis was analysed for both OA-HSFs (Figure 4) and MSCs (Figure 5) over 7 days. An early differentiation marker was also analysed to look at any changes specific to early stage osteogenic (RUNX2) or chondrogenic (SOX9) differentiation.

**Figure 4.**
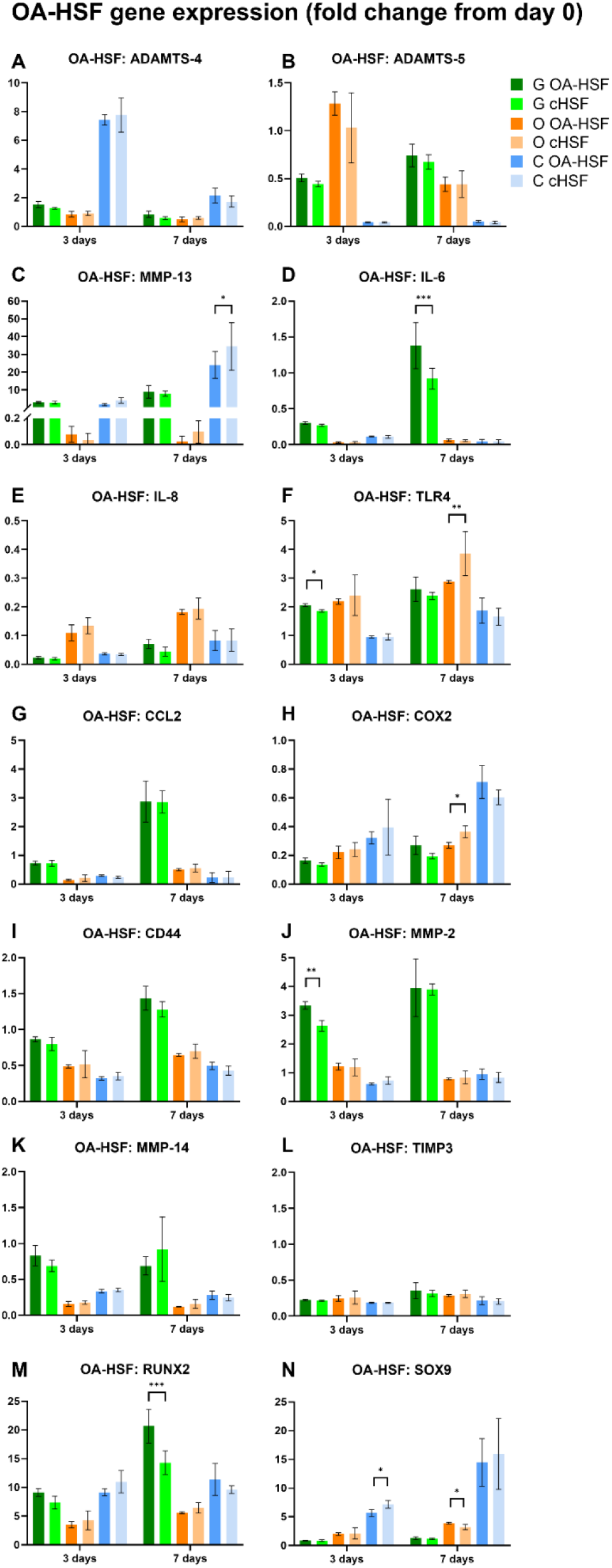
Gene expression of OA pro-inflammatory and pro-catabolic molecules, and early stage markers of osteogenic (RUNX2) and chondrogenic (SOX9) differentiation in OA-HSFs. OA-HSFs were co-cultured with fresh MSCs in growth (G), osteogenic (O), and chondrogenic (C) media for 3 and 7 days. The OA-HSFs were either pre-conditioned (cHSF) by pre-exposure to MSCs in co-culture for 3 days in growth medium, or not pre-conditioned (OA-HSF) which were grown by themselves for 3 days in growth medium, before being introduced to fresh MSCs. Expression levels were presented as fold change from MSCs at day 0, normalised to the housekeeping gene 18S. Data are represented as mean ± SD; N=4; *p < 0.05, **p < 0.01, ***p < 0.001.

**Figure 5.**
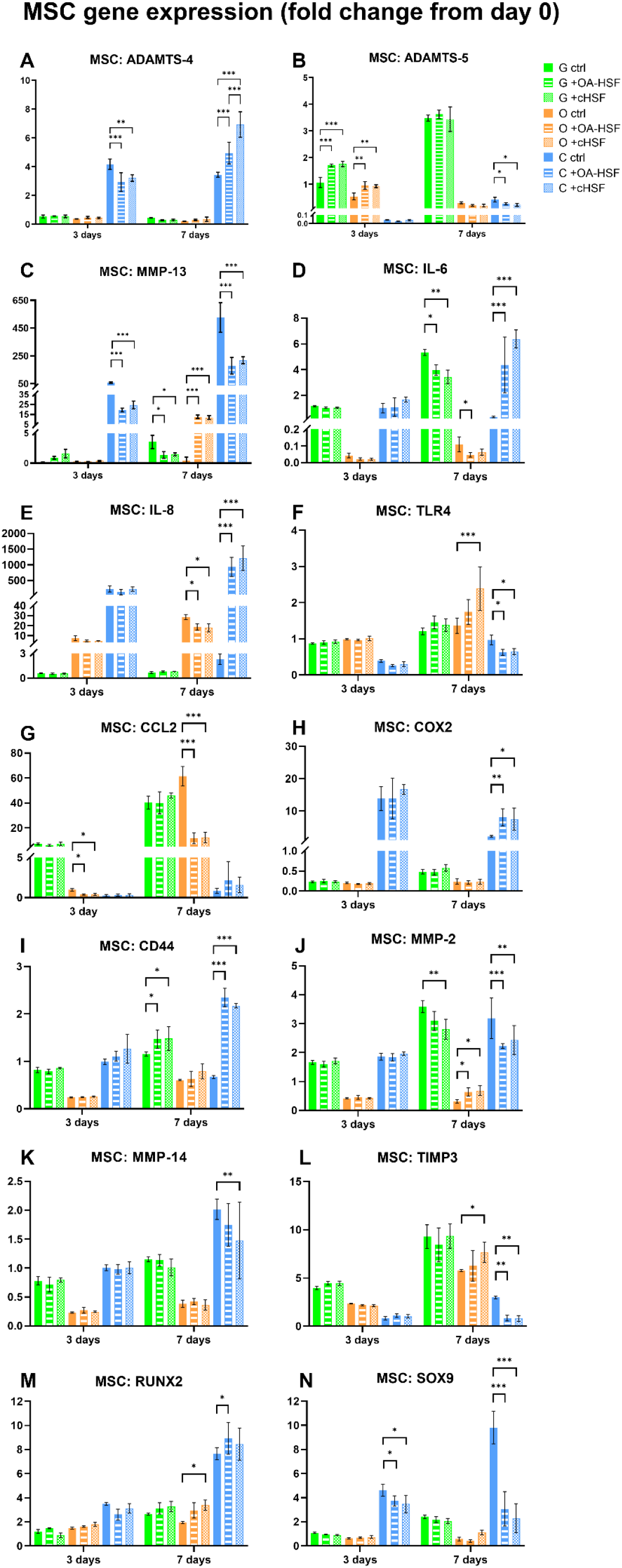
Gene expression of OA pro-inflammatory and pro-catabolic molecules, and early stage markers of osteogenic (RUNX2) and chondrogenic (SOX9) differentiation in MSCs. MSCs were grown by themselves or co-cultured with OA-HSFs in growth (G), osteogenic (O), and chondrogenic (C) media for 3 and 7 days. The OA-HSFs were either pre-conditioned (cHSF) by pre-exposure to MSCs in co-culture for 3 days in growth medium, or not pre-conditioned (OA-HSF) which were grown by themselves for 3 days in growth medium, before being introduced to fresh MSCs. Expression levels were presented as fold change from MSCs at day 0, normalised to the housekeeping gene 18S. Data are represented as mean ± SD; N=4; *p < 0.05, **p < 0.01, ***p < 0.001.

Pre-conditioning the OA-HSFs did not have significant effects on modifying their expression of OA-related pro-inflammatory and pro-catabolic molecules (Figure 4). Regardless of whether they were pre-exposed to MSCs or not, both groups of OA-HSFs co-cultured with fresh MSCs for 3 and 7 days showed similar expression levels for the vast majority of genes in all three media types and at both time points. There were some exceptions for OA-HSFs co-cultured with MSCs in growth medium, where pre-exposure to MSCs led to significant downregulation of IL-6, TLR4 and MMP-2 at 3 or 7 days. In chondrogenic medium, pre-conditioned OA-HSFs also showed significantly higher expression of the early chondrogenic marker SOX9 compared to control OA-HSFs at 3 days. These findings suggest that some of the paracrine conditioning effects of MSCs may be retained in the pre-conditioned OA-HSFs in the short term. However, there were also other exceptions where pre-conditioned OA-HSFs showed higher expression of some pro-inflammatory/catabolic factors, such as MMP-13, TLR4, and COX2 at 7 days, but exclusively in differentiation media which may be a consequence of global gene expression changes induced by early osteogenic or chondrogenic differentiation. Overall, *in vitro* pre-conditioning with MSCs did not induce significant changes in the pro-inflammatory and catabolic phenotype of OA-HSFs, suggesting that repeated MSC injections in an OA joint are not likely to induce sustained changes to this diseased environment.

Looking at MSCs which were co-cultured for 3 and 7 days with OA-HSFs that were pre-conditioned or not pre-conditioned, expression levels for the vast majority of genes were similar in all three media types. Significant differences in MSC gene expression in all media types were only seen when comparing the +OA-HSF or +cHSF group to the MSC control, but not between these two MSC groups co-cultured with OA-HSFs. The only exception was ADAMTS-4 at 7 days, where MSCs co-cultured with pre-conditioned OA-HSFs showed significantly higher expression of this gene compared to MSCs co-cultured with control OA-HSFs. For the osteogenic marker RUNX2, MSCs co-cultured with both OA-HSF groups showed higher expression at 7 days compared to the MSC control in osteogenic medium, possibly reflecting that early inflammatory priming is beneficial for kickstarting osteogenic induction. For the chondrogenic marker SOX9, MSCs co-cultured with both OA-HSF groups showed lower expression at 3 and 7 days compared to the MSC control in chondrogenic medium, reflecting the negative effects of an inflammatory environment on the induction of chondrogenic differentiation. For both of these early stage differentiation markers, there were no differences in expression in the MSC groups co-cultured with pre-conditioned or control OA-HSFs. Overall, these findings indicate that MSCs, whether co-cultured with OA-HSFs that were pre-conditioned or not pre-conditioned, do not change their expression levels of inflammatory or differentiation markers. In other words, the inhibitory effects of OA-HSFs on MSCs do not change, regardless of whether the OA-HSFs were previously exposed to MSC conditioning. Short-term exposure of osteoarthritic cells to MSCs may therefore be insufficient for sustained modifications to their diseased phenotype, and it may be unlikely that repeated MSC injections can improve therapeutic efficacy in OA.

## DISCUSSION

Through *in vitro* co-culture of human MSCs and OA-HSFs, the findings of this study suggested that despite the anti-inflammatory and trophic functions of MSCs, they could not provide long-term effects in correcting the osteoarthritic joint environment due to adopting the diseased behaviour of the surrounding cells. These findings have important implications for future OA therapies based on the use of stem cells. In addition to testing different cell sources, injection concentrations and administration frequencies in clinical trials, alternative approaches might be to work on correcting the catabolic environment within the osteoarthritic joint, or utilise the biological derivatives of stem cells which, unlike the living cell, will not respond in a negative way to the osteoarthritic environment.

In the first experiment, the MSCs were seen to provide a transient therapeutic effect on the OA-HSFs, resulting in immediate suppression of pro-inflammatory and pro-catabolic gene expression, however this trailed off at 7 days. This was consistent with the results of multiple clinical trials using intra-articular injections of MSCs for knee OA, which reported short-term improvement in most outcome measures post-administration but failure to maintain long-term therapeutic efficacy (Jiang *et al.*, 2021). It was also interesting to note that the short-term therapeutic effects of MSCs on OA-HSFs preferentially targeted genes directly involved in cartilage degradation, such as ADAMTS-4 and MMP-13, which respectively degrade aggrecan and type II collagen. These findings are consistent with clinical trials which reported short-term improvement of cartilage repair in knee OA treated with MSC injection (Ha *et al.*, 2019). It was previously hypothesised that the loss of therapeutic effect after MSC injection may be due to cell loss into the bloodstream or cell death at the defect site (Barry, 2019). The fact that our results demonstrated a decreased therapeutic effect of MSCs over time in co-culture with OA cells, where the MSCs remained localised and alive, suggest that the MSCs might be negatively responding to a diseased environment.

The above findings led us to perform the second experiment, to specifically investigate for the first time, the short- and long-term responses of MSCs in different culture environments relevant to an OA joint, when co-cultured with OA cells. It was interesting to find that in all cases where there were significant differences between groups cultured in growth medium, pro-inflammatory and pro-catabolic gene expression in MSCs was elevated when co-cultured with OA-HSFs at both short- and long-term time points. These findings suggest that MSCs may begin to display similar diseased and pro-inflammatory behaviour as the surrounding cells when placed in an osteoarthritic environment, limiting their beneficial effects. Ultimately, this altered MSC expression profile may even contribute to ongoing and worsening matrix degradation and progression of OA. Intriguingly, there were a number of cases in long-term (21 day) co-culture where MSCs exposed to OA-HSFs downregulated inflammatory marker expression, but exclusively in osteogenic or chondrogenic medium. This may suggest an exciting prospect for long-term conditioning of the OA joint to an environment that induces bone or cartilage regeneration to promote the paracrine functions of MSCs. Further studies are needed to explore the feasibility of this approach.

It was interesting to explore the influence of OA cells on the differentiation capacity of MSCs in osteogenic and chondrogenic induction media, which is one of the defining characteristics of MSCs (Chamberlain et al., 2007). MSCs exposed to OA-HSFs in long-term co-culture showed significantly impaired ability to undergo osteogenic and chondrogenic differentiation, notably through the markedly reduced expression of mid-late stage differentiation markers and prominent loss of staining area in histological analysis. The closest comparison to our results comes from a study that examined gene expression in synovial fluid-derived MSCs from patients with advanced knee OA (Sanjurjo-Rodriguez et al., 2020). These MSCs showed a decrease in osteogenic gene expression, specifically BSP and RUNX2, compared to control MSCs from subchondral bone. Interestingly, this study also described a 5 to 50-fold increase in the expression of pro-catabolic proteins including MMP1 and ADAMTS-5 in the OA synovial fluid MSCs. Overall, our results from the second experiment suggest that the MSCs can lose their restorative and trophic characteristics under the influence of diseased OA cells, which may contribute to their short-lived therapeutic effects as observed in clinical trials (Jiang *et al.*, 2021).

Some efforts have been made to unearth the mechanisms by which the secretion of paracrine factors in MSCs is affected by inflammatory conditioning. One study used proteomic analysis to identify 118 factors secreted by human adipose-derived MSCs in response to TNF-α exposure, and found that several chemokines were secreted at significantly higher levels compared to quiescent MSCs, including IL-6, IL-8, CXCL6, and MCP-1 (Lee *et al.*, 2010). Since TNF-α is one of the key inflammatory cytokines and may be linked to OA progression, these findings implicate that the paracrine functions of MSCs can be converted to a pro-inflammatory state in an OA environment, complementing the results of our study. In the only other study that established *in vitro* 3D models of OA to study MSC interactions, the OA models were tuned into high and low inflammatory environments (Diaz-Rodriguez *et al.*, 2019). In a high-inflammatory OA environment, MSCs induced osteoarthritic chondrocytes to produce markedly lower levels of IL-1β, interferon (IFN)-γ, MMP-9, and MMP-13, and reduced macrophage activation, which were different to the responses seen in a low-inflammatory OA environment. Together with the findings of our study, there is emerging evidence to suggest that the immunomodulatory and therapeutic effects of MSCs are influenced by the OA environment, and may be further dependent on the severity of joint inflammation.

The third experiment could be considered to simulate repeated administration of MSCs to an OA joint. If repeated exposure of OA cells to MSCs could induce a greater beneficial effect, we would expect differences in inflammatory gene expression between the control and pre-conditioned OA-HSF groups, for both the OA-HSFs and also the MSCs co-cultured with OA-HSFs. However, overall trends suggested that re-exposure of OA-HSFs to MSCs, even when conducted over a short time frame of a few days, was not sufficient to induce sustained changes to their diseased phenotype. In the OA-HSFs, there were some transient effects where MSC pre-conditioning led to downregulation of certain inflammatory markers in growth medium, which perhaps justifies some of the increased therapeutic benefits seen with repeated MSC injections for knee OA in clinical trials (Matas *et al.*, 2019; Song et al., 2018). No significant differences in gene expression were observed for MSCs co-cultured with OA-HSFs, whether pre-conditioned or not, suggesting that repeated exposure of OA-HSFs to MSCs does not reduce their negative influence on the paracrine activities of MSCs. These findings may implicate that single or even multiple MSC injections will not lead to a sustained therapeutic effect in OA joints.

Difficulties are encountered when interpreting the outcomes of published clinical studies reporting the use of MSC injections to treat OA, due to large variations in the tissue sources of MSCs, isolation and expansion protocols, injection doses and frequencies, delivery vehicles, patient cohorts, outcome measures, length of follow-up, and so on (Ha *et al.*, 2019; Jiang *et al.*, 2021; McIntyre et al., 2017). The number of patients recruited to the majority of existing clinical studies is also relatively small. For instance, two trials testing repeated administration of MSCs derived from umbilical cord (Matas *et al.*, 2019) and adipose tissue (Song *et al.*, 2018) respectively, each involved 6 to 9 patients per treatment group. In both trials, improvements were seen in outcome measures related to pain (such as VAS and WOMAC) and quality of life (such as SF-36). However, when compared to repeated injections of hyaluronic acid, there was no improvement in MRI score as an objective measure of cartilage repair at 12 months (Matas *et al.*, 2019). The other study compared low, medium, and high doses of MSCs (10-50 million cells) all administered with 3 injections rather than between single and multiple injections (Song *et al.*, 2018). Although cartilage volume increased over the follow-up period, this had returned to baseline by 96 weeks. The findings of our study complement the existing clinical evidence on multiple MSC injections for knee OA. There are also interesting contradictions regarding the optimal dose of MSCs in the clinical treatment of knee OA. Some studies show the greatest benefits at the highest dose, ranging from 50 (Song *et al.*, 2018) to 100 million cells (Jo et al., 2017), although this may also give rise to additional risks of pain, joint infection and swelling (Gupta et al., 2016). Others have reported greater therapeutic benefits of low-dose MSC injections, where 10 million bone marrow-derived MSCs outperformed 100 million (Lamo-Espinosa et al., 2016), or 2 million adipose-derived MSCs outperformed 10 and 50 million (Pers et al., 2016). The array of inconsistent clinical results raises uncertainty for the long-term benefits of periodic MSC treatment for OA.

By investigating the reciprocal responses of MSCs and OA cells using an *in vitro* co-culture model, our study provided novel insights into how the paracrine functions of MSCs might be affected by the OA environment, and the implications of this on the long-term therapeutic efficacy of single or multiple MSC injections for clinical OA treatment. It is important to keep in mind when interpreting the results of our study that the experiments were conducted *in vitro* using isolated cell types in controlled biochemical environments, and may not recapitulate the complex *in vivo* milieu of an OA joint. Our study also alludes to an exciting prospect of improving the therapeutic efficacy of MSCs through priming techniques, such as by exposing the MSCs to hypoxia (Pattappa et al., 2020) or compounds including vitamin E (Bhatti et al., 2017), which have led to improved treatment effects in OA *in vivo* models.

## EXPERIMENTAL PROCEDURES

### Cell sources

OA-HSFs were isolated from the synovium of patients with knee OA undergoing joint replacement surgery at Royal North Shore Hospital and North Shore Private Hospital, as previously described (Smith *et al.*, 2013). Briefly, tissue normally discarded as part of standard surgical care was collected with written informed consent from OA patients undergoing total or partial knee replacement surgery (Northern Sydney Central Coast Health HREC Protocol 0902-034M). De-identified tissues were provided to the laboratory and synovial fibroblasts (HSF) were isolated by proteolytic digestion, grown to confluence in a T75 flask in DMEM supplemented with 10% (v/v) foetal bovine serum (FBS) and 2mM glutamine, trypsinised and cryopreserved frozen in aliquots (1 × 10^7^ cells/mL) (Melrose et al., 2003; Smith and Ghosh, 1987). In the present study, OA-HSFs from 2 patients (69 and 73 year old males, undergoing total knee replacement) were pooled and used at the 6^th^ to 10^th^ passage for all experiments.

Human bone marrow-derived MSCs were purchased from RoosterBio (MSC-003; Frederick, Maryland, USA). Before use in experiments, MSCs were expanded in RoosterNourish™-MSC (RoosterBio) according to the manufacturer’s instructions. Briefly, MSCs were thawed and resuspended in RoosterNourish™-MSC, and plated in T-75 flasks at an average seeding density of 3.3×10^3^ cells per cm^2^. The MSCs were grown until 70-80% confluency before use. MSCs used for experiments were at passage three, and their trilineage differentiation ability was confirmed by adipogenic, chondrogenic, and osteogenic assays before use.

### Co-culture experiments

MSCs were co-cultured with OA-HSFs in three types of culture media, simulating the types of conditions relevant for OA: (1) Growth medium: Dulbecco’s Modified Eagle Medium (DMEM) supplemented with 10% (v/v) foetal bovine serum (FBS) and 2 mM L-glutamine; (2) Osteogenic medium: StemPro^TM^ Osteogenesis Differentiation Kit from Life Technologies (Grand Island, NY, USA); (3) Chondrogenic medium: StemMACS^TM^ ChondroDiff Medium from Miltenyi Biotec (Bergisch Gladbach, Germany). MSCs were cultured in 12-well plates, seeded at a density of 4×10^4^ cells per well in monolayer for growth and osteogenic cultures, or seeded as a micromass containing 3×10^5^ cells for chondrogenic cultures. For co-culture with OA-HSFs, the HSFs were seeded at a density of 2×10^4^ cells per insert in 12-well Transwell^®^ inserts with polycarbonate membrane and pore width of 0.4 μm (Corning, New York, US). Inserts with seeded HSFs were then placed in 12-well plates with growth, osteogenic, or chondrogenic MSC cultures to establish the *in vitro* co-culture model. The permeable inserts allowed exchange of culture medium and metabolites between MSCs and OA-HSFs, but not physical contact between them. All cultures were incubated at 37°C in 5% CO_2_, and 50% of the culture medium was replaced three times per week.

Three studies were conducted. (1) To confirm the short-term effects of MSCs on OA cells, OA-HSFs were cultured with (HSF + MSC) or without (HSF control) MSCs in growth medium for 3 and 7 days. (2) To test whether OA cells can modify the behaviour of MSCs in the short and longer term, MSCs were cultured with (MSC + HSF) or without (MSC control) OA-HSFs in growth, osteogenic, and chondrogenic media for up to 21 days. (3) To test whether exposing OA cells to MSCs can cause sustained changes in their behaviour and have positive effects on tissue repair, OA-HSFs were either pre-conditioned by first co-culturing with MSCs for 3 days (cHSF), or not pre-conditioned and simply cultured in growth medium for 3 days (HSF), and both groups were subsequently co-cultured with fresh MSCs in growth, osteogenic, and chondrogenic media for 3 and 7 days.

### Gene expression analysis by quantitative RT-PCR

MSCs and OA-HSFs were harvested separately from each co-culture assay at the indicated time points for gene expression analysis. RNA was extracted from MSCs and OA-HSFs using the Isolate II RNA Micro Kit (Bioline, Canada) following the manufacturer’s instructions. MSC chondrogenic micromasses were first snap-frozen in liquid nitrogen and minced before placing in the lysis buffer, while lysis buffer was added directly to all monolayer MSCs or OA-HSFs in the well or Transwell^®^ insert. RNA concentration of each sample was quantitated using Nanodrop 2000 (Thermo Fisher, Australia), and reverse transcription to cDNA (Omniscript, Qiagen) was conducted using 500 ng RNA per sample, using random pentadecamers (50 ng/mL, Sigma-Genosys) and RNase inhibitor (10 U per reaction, Bioline). The resulting cDNA was subjected to real-time PCR in a Rotorgene 6000 (Corbett Life Science, New South Wales, Australia) using Immomix (2× dilution; Bioline), SYBR Green I (10,000× dilution; Cambrex Bioscience) and 0.3 μM human-specific primers (Sigma-Aldrich), as shown in Supplementary Table S1. The expression levels of genes were normalised to those for the 18s housekeeping gene using the comparative Ct (2^−ΔΔCt^) method, and results were expressed as fold changes in gene expression relative to unstimulated control cells at day 0 (before the commencement of cultures). Pro-inflammatory and pro-catabolic response in the MSCs and OA-HSFs were evaluated using expression of OA-related genes (IL-6, IL-8, MMP-2, MMP-13, ADAMTS4, ADAMTS5, CD44, TLR4, COX-2, CCL2), and differentiation in the MSCs was evaluated using gene markers for osteogenesis (RUNX2, BSP, SPP1) and chondrogenesis (SOX9, COL2A1, ACAN).

### Histological analysis

MSC differentiation was additionally evaluated by histology at 21 days. For osteogenic cultures, monolayer MSCs were fixed in 10% neutral buffered formalin (NBF) for 15 minutes, washed, and stained with Alizarin Red S to visualise calcium deposition. For chondrogenic cultures, MSC micromasses were fixed in 10% NBF for 4 hours, then embedded in paraffin, sectioned and stained with toluidine blue to visualise proteoglycan deposition. Stained samples were visualised using light microscopy.

### Statistical analysis

Data for all experiments were obtained from four independent samples, except for histological analyses which were obtained from two independent samples. Gene expression data were expressed as mean ± standard deviation (SD). Statistical analyses were performed using GraphPad Prism V9.4.0 (GraphPad Software, USA). Differences in gene expression levels between groups were analysed using two-way ANOVA, followed by Sidak’s multiple comparisons test, with p < 0.05 considered statistically significant.

## Author contributions

J.J.L. and C.B.L. conceptualised the study and designed the experiments. J.J.L. conducted the experiments. V.S., J.L. and J.J.L. performed data analysis and wrote the paper. All authors critically revised the paper.

## Acknowledgements

The authors acknowledge Ms Susan Smith for the preparation of histological samples, and funding from the National Health and Medical Research Council (GNT1120249), Arthritis Australia, and The Lincoln Centre.

## Declaration of interests

The authors declare no competing interests.

## Notes

### Competing Interest Statement

The authors have declared no competing interest.

