## Supplementary material for "Understanding the effects of mesenchymal stromal cell therapy for treating osteoarthritis using an *in vitro* co-culture model": Table S1

**Supplementary Table S1.** Primer sequences for genes evaluated by real-time RT-PCR

| Gene | 5' to 3' forward sequence | 5' to 3' reverse sequence |
|---|---|---|
| 18S | ACC ATA AAC GAT GCC GAC CG | CAA TCT GTC AAT CCT GTC CGT GTC |
| ADAMTS-4 | AGA CAC AGG CAG GGA GAG ACA AAG | GGA GAA AAC TTA GTC CTT GGG CTT G |
| ADAMTS-5 | AAC TCC CAG GAC AGA CCT ACG ATG | GCA GAT TCT CCC CTT TCC ACA AG |
| CCL2 | GCT GAG ACT AAC CCA GAA ACA TCC | AAT GAA GGT GGC TGC TAT GAG C |
| CD44 | AAA GGA GCA GCA CTT CAG GA | TGT GTC TTG GTC TCT GGT AGC |
| COX2 | GAC AGT CCA CCA ACT TAC AAT GCT G | GCT GCT TTT TAC CTT TGA CAC CC |
| IL-6 | GAA GAT TCC AAA GAT GTA GCC GC | GAA GGT TCA GGT TGT TTT CTG CC |
| IL-8 | AAC TTT CAG AGA CAG CAG AGC ACA C | CAC AGT GAG ATG GTT CCT TCC G |
| MMP-2 | TGA CGG AAA GAT GTG GTG TG | CTC CTG AAT GCC CTT GAT GT |
| MMP-13 | TGA CGG AAA GAT GTG GTG TG | CTC CTG AAT GCC CTT GAT GT |
| MMP-14 | CCA GGG TCT CAA ATG GCA ACA TAA TGA AA | CCA TGG AAG CCC TCG GCA AA |
| TIMP3 | TGC CCT TCT CCT CCA ATA CA | CTT CCT TCC CTC CCT CAC TC |
| TLR4 | GCT TTC ACT TCC TCT CAC CCT TTA G | CTG GCA TCA TCC TCA CTG CTT C |
| RUNX2 | CCA GAT GGG ACT GTG GTT ACT GTC | CTGG GGA GGA TTT GTG AAG ACG |
| BSP | ATG GCC TGT GCT TTC TCA ATG | GGA TAA AAG TAG GCA TGC TTG |
| OP | ACG CCG ACC AAG GAA AAC TC | GGA GAT TCT GCT TCT GAG ATG GG |
| OC | ATG AGA GCC CTC ACA CTC CTC G | GTC AGC CAA CTC GTC ACA GTC C |
| SOX9 | AAA CGG TGC TGC TGG GAA AC | CTC CTT TGC TTG CCT TTT ACC TC |
| COL2A1 | CAG TTC GGA CTT TTC TCC CCT C | AGT TTC CTG CCT CTG CCT TGA C |
| ACAN | TCA CCA TCC CCT GCT ATT TCA TC | TCT CCT TGG ACA CAC GGC TC |
| COL1A1 | ACA GGG CGA CAG AGG CAT AAA G | AAC AGG ACC AGC ATC ACC AGT G |
